# A male-derived volatile sex pheromone in *Caenorhabditis* nematodes identified through its mimicry by a predator

**DOI:** 10.1101/2025.09.12.675966

**Authors:** Matthew R. Gronquist, Xuan Wan, Daniel Leighton, Yuki Togawa, Marika Sagawa, Paul W. Sternberg, Frank C. Schroeder, Ryoji Shinya

## Abstract

Recent studies demonstrated that the predacious fungi *Arthrobotrys oligospora* emits a mixture of volatile chemical cues that function to attract nematode prey. The strong attraction elicited by one of the mixture components, methyl 3-methyl-2-butenoate (MMB), was highly female- and hermaphrodite-specific within several *Caenorhabditis* species, including *C. remanei* and *C. elegans*, suggesting that MMB might function as a mimic of an endogenous, male-produced, volatile sex pheromone (VSP) within these species. Here, we report evidence that MMB is produced by *C. remanei* males at levels that are attractive to *C. remanei* females and *C. elegans* hermaphrodites. Notably, MMB production was not detected for *C. elegans* males; a finding which correlates with behavioral assays for which worm-conditioned media (WCM) prepared from *C. remanei*, but not from *C. elegans* adult males is strongly attractive to both *C. remanei* females and *C. elegans* hermaphrodites. Our findings establish MMB as the first chemically identified VSP in nematodes and show that *A. oligospora* exploits a dual strategy of chemical deception—mimicry and eavesdropping—to enhance prey capture.

## Introduction

Chemical mimicry, wherein an organism produces or exploits chemical signals closely resembling the communication cues of another species, provides fascinating examples of interspecific interactions and evolutionary adaptations^1,2^. Within predator-prey and parasite-host systems, chemical mimicry often involves the exploitation of pheromone-mediated communication, enabling the predator or parasite to deceive prey or hosts. Examples of such aggressive chemical mimicry include adult female bolas spiders that lure male moths by mimicking their sex pheromones^3,4^, and slave-making ants that produce alarm pheromone mimics to disrupt the social structure of host colonies^5^. Hosts may, however, evolve the ability to detect and respond to parasitic chemical cues. For example, Ascr#18, a conserved ascaroside signalling molecule produced by some plant-parasitic nematodes, triggers broad-spectrum pathogen resistance in a variety of host plants^6^. Mutualistic systems may furthermore use mimicry for reciprocal benefit. The plant-parasitic pinewood nematode secretes ascaroside pheromones that induce metamorphosis in its vector beetle, while beetle-derived ascarosides attract nematode larvae, facilitating dispersal to new host trees^7^. These dynamics highlight how chemical mimicry evolves not only as a tool for deception but also as a mediator of cooperative strategies, blurring traditional boundaries between parasitism and mutualism.

Recent studies of the nematode-trapping fungus *Arthrobotrys oligospora* revealed a remarkable instance of chemical mimicry wherein the fungus secretes a mixture of volatile compounds, including those representing olfactory cues of food, to lure prey nematodes^8^. One component of the fungal-derived volatile mixture, methyl 3-methyl-2-butenoate (MMB), stood out due to its particularly strong attraction that was specifically directed toward females and hermaphrodites of several *Caenorhabditis* species. This female- and hermaphrodite-specific attraction suggested that MMB might function as a mimic of a yet unidentified male-produced volatile sex pheromone (VSP) within these species. Indeed, in the dioecious species *C. remanei*, females exhibited strong attraction to MMB, whereas males were repelled by it.

In the androdioecious species, *C. elegans*, hermaphrodites showed strong attraction, and males displayed only weak attraction. The existence of female- and hermaphrodite-derived VSPs which are attractive to males is well documented within *C. elegans* and related species^9–12^, although the specific chemical signals involved remain unknown. By contrast, evidence for the existence of a male-derived VSP that is attractive to females or hermaphrodites has been less clear. Chasnov et al. had previously found WCM obtained from *C. remanei* males to be weakly attractive to both *C. remanei* females and *C. elegans* hermaphrodites^9^. In the case of *A. oligospora*, whether MMB itself might represent a chemical mimic of an endogenous male-produced VSP within *Caenorhabditis* nematodes was unknown. With this work we provide evidence that MMB is indeed a volatile sex pheromone produced by adult males, but not females, of *C. remanei*. Notably, we find no evidence of MMB production within *C. elegans* males or hermaphrodites, despite the attraction that MMB also elicits toward *C. elegans* hermaphrodites, highlighting an intriguing difference between these species that is potentially linked to their distinct reproductive modes, dioecy and androdioecy.

We used solid phase microextraction (SPME)^13^ to sample headspace volatiles present above live cultures of *C. remanei* and *C. elegans* adult males, as well as those of females and hermaphrodites from both species, respectively. Analyses by comprehensive two-dimensional gas chromatography-mass spectrometry (GC x GC-MS) and by gas chromatography-mass-spectrometry (GC-MS) revealed MMB to be present in samples obtained from *C. remanei* males but not those obtained from females, nor from samples collected from *C. elegans* males or hermaphrodites. MMB was present in the *C. remanei* adult male cultures at concentrations that were found to be strongly attractive to *C. remanei* females in behavioral assays. Our finding that MMB is produced by *C. remanei*, but not *C. elegans*, males additionally correlates with further assays for which WCM prepared from *C. remanei* adult males was strongly attractive to both *C. remanei* females and *C. elegans* hermaphrodites, whereas WCM prepared from *C. elegans* adult males showed no meaningful attraction to either.

## Results

### MMB is produced by *C. remanei* adult males

SPME headspace sampling of cultures of *C. remanei* adult males prepared in sealed vials, followed by GC x GC-MS and GC-MS analyses revealed a peak which was unique to the *C. remanei* male samples and whose electron-impact mass-spectrum and chromatographic retention time were indistinguishable from those obtained for an authentic sample of MMB (Fig. 1). The identification of potential pheromone candidates by gas-chromatographic mass-spectral analyses is often complicated by the occurrence of numerous background peaks, as well as other cryptic or irrelevant peaks. In the present case, however, the identification of MMB was immediate, since we had recently encountered this identical compound as a potential female-and hermaphrodite-attractive pheromone mimic produced by the predacious fungi *A. oligospora*^8^. MMB was not detected for any SPME samples collected from adult *C. remanei* females, nor from any *C. elegans* sample, including *C. elegans* males (Fig. 2A).

**Fig. 1:**
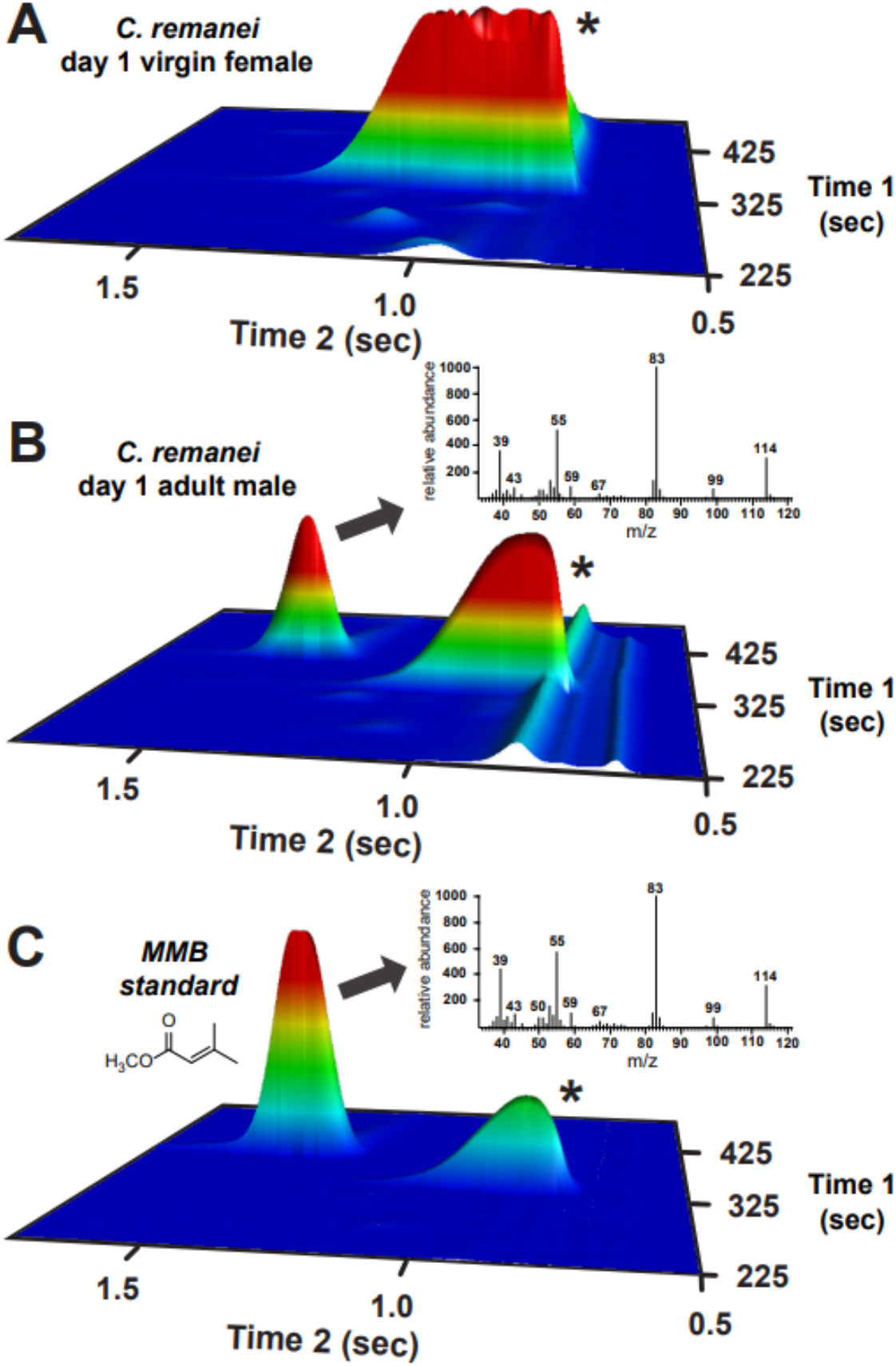
MMB is produced specifically by *C. remanei* adult males. Representative GC x GC-MS chromatograms (oblique view) and mass spectra obtained for SPME headspace sampling of *C. remanei* males and females, as well as for an authentic sample of MMB. A peak with retention time and mass spectrum (insets) matching those of methyl 3-methyl-2-butenoate (MMB) was prominent in the *C. remanei* male samples, but was not detected in those from *C. remanei* females. **(A)** *C. remanei* 1-day-old virgin females. **(B)** *C. remanei* 1-day-old adult males. **(C)** Comparative analysis of an authentic sample of MMB. The chromatograms are scaled uniformly and the retention time regions shown are selected to clearly show relevant peaks. The second peak which is present in each chromatogram (marked with an asterisk) is a persistent, background peak likely arising as an artefact from the SPME fiber. Additional complete chromatograms and mass spectra, including those for *C. elegans* may be found in the supplementary information.

**Fig. 2:**
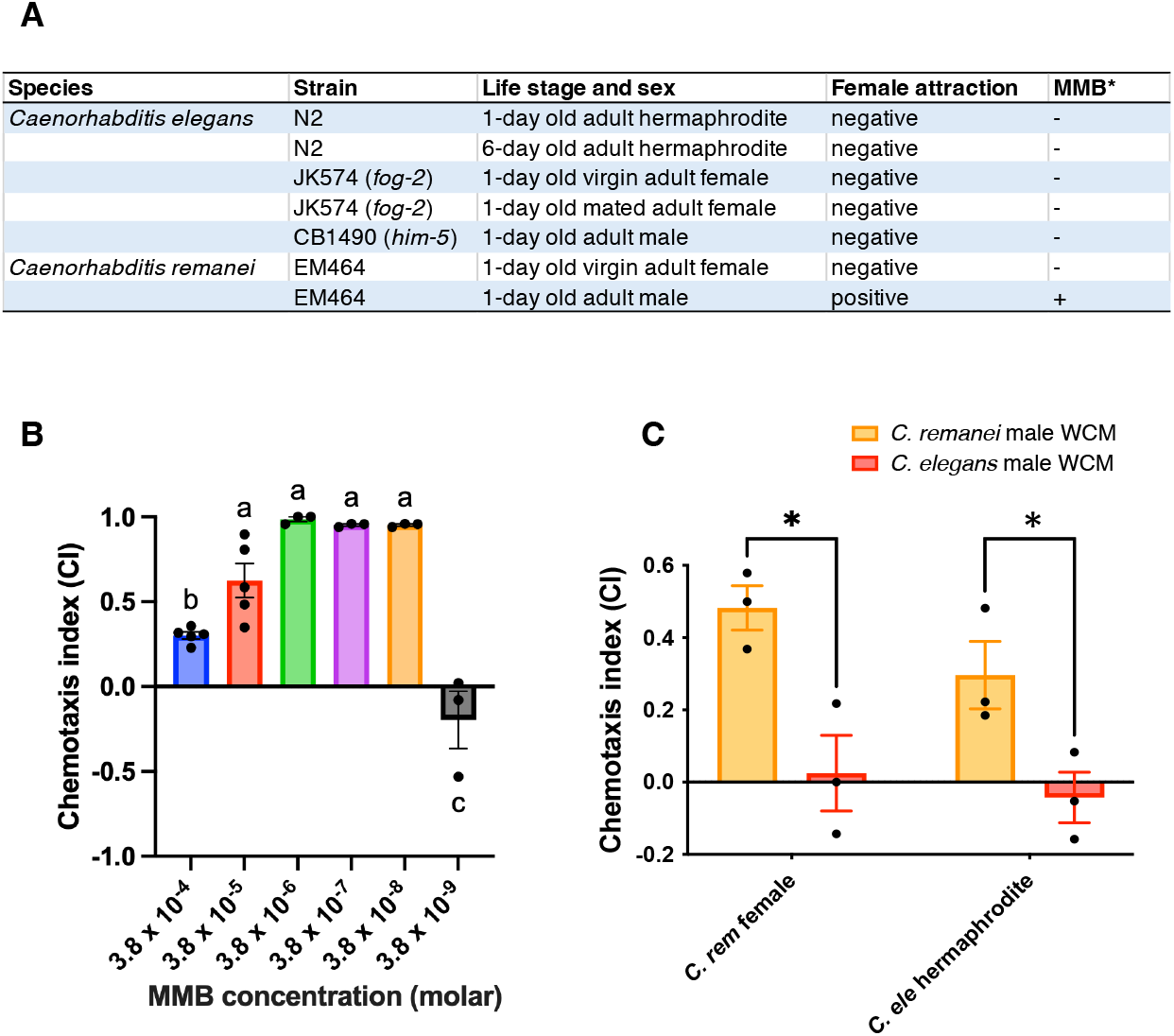
**(A)** Detection of MMB in *C. remanei* and *C. elegans* across different sexes and developmental stages. **(B)** Chemotaxis responses of *C. remanei* females to MMB, ranging from 3.8 × 10^-4^ M to 3.8 × 10^-9^ M. Different letters indicate statistically significant differences among concentrations according to Tukey’s multiple comparison test (p < 0.01). **(C)** Chemotaxis responses of *C. remanei* females and *C. elegans* hermaphrodites to WCM extracted from *C. remanei* and *C. elegans* males. Statistical significance was determined using a two-tailed t-test (p < 0.05). Asterisks indicate significant differences. Error bars indicate standard error.

### Methyl 3-methyl-2-butenoate (MMB) is a pheromone of *C. remanei* males

Hsueh et al. found that MMB is produced by the nematode-trapping fungus *A*.*oligospora* and that it triggers strong adult hermaphrodite and female-specific attraction in *Caenorhabditis* nematodes^8^. Having established that MMB is additionally produced by males of *C. remanei*, we used a chemotaxis assay optimized for volatile sex pheromones, as originally developed by Leighton et al.^10^, to evaluate the concentration dependence of MMB attraction for *C. remanei* females. While Hsueh et al. demonstrated attraction to MMB by *C. elegans* hermaphrodites over a broad concentration range, their assay tested only a single concentration for *C. remanei* females^8^. We therefore performed assays using a dilution series of MMB and found that *C. remanei* females are strongly attracted to MMB within the concentration range of 3.8 × 10^−6^-to 3.8 × 10^−8^-M (Fig. 2B). Quantitative analyses by SPME-GC-MS determined the concentration of MMB within the *C. remanei* adult male samples to be on the order of 10^-7^ M, notably within the observed attractive concentration range.

Chasnov et al. had previously found WCM obtained from *C. remanei* males to be weakly attractive to both *C. remanei* females and *C. elegans* hermaphrodites, whereas WCM obtained from *C. elegans* males showed no meaningful attraction to either species^9^. We carried out additional assays to evaluate the attractiveness of WCM prepared from *C. remanei* and *C. elegans* males with respect to the disparate production of MMB observed within these species. We found significantly strong attraction for *C. remanei* females (CI = 0.48, *p < 0*.*05, t-*test) and *C. elegans* hermaphrodites (CI = 0.30, *p < 0*.*05, t-*test) (Fig. 2C) to WCM prepared from *C. remanei* adult males. No significant attraction for *C. remanei* females (CI = 0.02) or *C. elegans* hermaphrodites (CI = -0.04) was observed for WCM prepared from *C. elegans* adult males.

## Discussion

With this work, we have identified MMB as a female-attractive, VSP produced by males of

*C. remanei*. By classical definitions, sex pheromones are chemical signals released by one sex that elicit mating-related behaviors—typically attraction—in the opposite sex^14^. MMB fits these criteria: it is produced specifically by adult *C. remanei* males and induces strong chemotactic attraction in conspecific females. In contrast to previously identified nematode sex pheromones, such as ascarosides and vanillic acid—the latter produced by the plant-parasitic nematode *Heterodera glycines*,^15^ which are water-soluble and lack significant volatility, MMB is volatile and the elicited attraction occurs through the air over greater distances, independent of diffusion within the growth medium. To our knowledge, MMB represents the first VSP enabling airborne communication between sexes to be chemically identified in nematodes. We find that MMB triggers sex-specific attraction in *C. remanei* females at concentrations as low as 3.8 × 10^-8^ M as a single component (Fig. 2C), a threshold that is notably lower than the working concentrations typically required for individual pheromone constituents to elicit comparable responses in multi-component blends.

Although MMB attracts females or hermaphrodites from both *C. remanei* and *C. elegans*, respectively, it was not detected in any *C. elegans* samples (Fig.2A). As *C. elegans* evolved from a dioecious to an androdioecious species, the ability of males to produce MMB could plausibly have decayed due to a lack of selective pressure, whereas *C. elegans* hermaphrodites retained the ability to respond to MMB, possibly because MMB has a role other than that of a sex pheromone in *C. elegans*, e.g., as a food cue. The possibility that *C. elegans* males might produce MMB under specific, but as yet unknown, conditions furthermore exists. Analyses of the pheromone systems of additional *Caenorhabditis* species may help clarify MMB function in *C. elegans*.

The structure of MMB provides additional insight into its possible biosynthetic origin and ecological relevance. Similar to many insect pheromones^16^, the chemical structure of MMB suggests that it originates from metabolism of essential amino acids. Leucine degradation proceeds via the coenzyme-A derivative of 3-methyl-2-butenoate^17^, which following hydrolysis and methylation would yield MMB. This biosynthetic origin of MMB from leucine would be consistent with the notion that pheromones can represent fitness signals, e.g., indicating sufficient access to food sources containing essential amino acids^18^. For example, the biogenesis of a *C. remanei* female-produced, male-attractant ascaroside pheromone depends on cyclopropyl fatty acid synthase (cfa), which is expressed in bacteria during their stationary growth phase, linking pheromone production to microbial metabolic cues^19^. Notably, MMB bears structural similarity to the methyl ester of tiglic acid, a motif that also appears in known *Caenorhabditis* metabolites. For example, the tigloyl moiety is incorporated into certain ascarosides^20^, suggesting possible chemical convergence or functional overlap between distinct classes of signaling molecules.

MMB was previously identified as a suspected pheromone mimic produced by the nematode-trapping fungus, *A. oligospora*^8^. Remarkably, MMB is not the only nematode chemical cue to have been compromised by *A. oligospora*. Multiple species of nematode-trapping fungi, including *A. oligospora*, have been shown to detect nematode-derived ascarosides as a cue to the presence of nearby prey, responding by initiating trap formation^21^. By emitting MMB, the fungus mimics male *C. remanei* reproductive signals, targeting adult females via a high-fidelity cue tied to a reproductive behavioral strategy, thereby optimizing energy investment in trap formation. The cross-species attractiveness of MMB toward females and hermaphrodites within *Caenorhabditis* nematodes may have furthermore made MMB a particularly effective target for the basis of this deception. Future study of whether fungal MMB production is dynamically regulated in response to prey density or mating behavior, mirroring adaptive strategies seen in other predator-prey systems will be of interest^7^. Such plasticity would further highlight the role of chemical signaling networks as battlegrounds for evolutionary innovation.

The present work demonstrates that *A. oligospora* is able to exploit two distinct communication networks operating within its nematode prey, using two distinct strategies of chemical deception: eavesdropping and mimicry^1,2^. Thus, *A. oligospora* deploys MMB as a chemical mimic to attract prey while also eavesdropping on ascaroside signaling to detect the presence of prey^21^. These dual strategies illustrate the remarkable sophistication of predator-prey interactions mediated by chemical signaling and underscore the ecological importance of volatile communication in soil-dwelling nematodes. Unlike mutualistic systems where ascaroside exchange drives cooperative dispersal (e.g., pinewood nematode-beetle interactions)^7^, this antagonistic interaction underscores chemical mimicry’s dual role in mediating both exploitation and mutualism.

## Methods

### Nematode preparation for solid-phase microextraction (SPME) sampling in *C. elegans* and *C. remanei*

N2, JK574 (*fog-2*) and CB1490 (*him-5*) strains of *C. elegans* and the EM464 strain of *C. remanei* were used for headspace sampling. The nematodes were propagated on NGM plates that had been seeded with *Escherichia coli* OP50 at 20°C. The nematodes were synchronized by bleaching and at the fourth larval stage (L4) the hermaphrodites/females and males were sorted and transferred onto new OP50 plates. The following day, 1000 each of the nematodes (*C. elegans* young adult N2 hermaphrodites, *fog-2* females, *him-5* males or *C. remanei* EM464 males) were picked into 1 ml of M9 buffer in 4 ml clear glass vials which were sealed with PTFE/silicone septa and immediately subjected to SPME sampling. Sperm depleted N2 hermaphrodites were prepared by transferring to a fresh OP50 plate every day until no more eggs were laid, at which point 250 of the sperm-depleted, old hermaphrodites in 250 µl of M9 buffer were subjected to headspace sampling as described above. Mated *fog-2* females were prepared by overnight incubation with 1-3 of N2 males on an OP50 plate, following which 250 mated females with confirmed egg laying were used for the headspace sampling.

### Headspace sampling of volatiles

SPME was used to sample volatiles present in the headspace above worm cultures prepared in sealed 4 mL vials as described above. All SPME fibers (divinylbenzene/carboxen/polydimethylsiloxane, 50/30 mm; Supelco, Bellefonte, PA) were pre-conditioned for 10 min in the GC injector (250°C) prior to use. For a typical collection, each fiber was inserted through the vial septa and exposed for a time period of 16 hr at room temperature. Fibers were either immediately analyzed or sealed in SPME fiber assembly storage devices (Supelco) to await analysis.

For quantitative analyses, standard dilutions of MMB prepared as 1 mL aliquots in sealed 4 mL vials were sampled by SPME for 16 hrs and immediately analyzed by GC-MS. A two-point calibration curve prepared from those samples which most closely bracketed the MMB peak areas observed in the *C. remanei* male samples was used to determine the MMB concentration for those samples.

### Gas Chromatographic Mass Spectral Analyses

Analyses of SPME samples were carried out using both comprehensive two-dimensional gas chromatographic mass spectrometry (GC x GC-MS), as well as one-dimensional gas chromatography mass-spectrometry (GC-MS). Methyl 3-methyl-2-butenoate (MMB) was positively identified based on a comparison of its electron-impact mass spectrum and gas-chromatographic retention time with those of an authentic sample purchased from Sigma Aldrich.

### GC x GC-MS analyses

Comprehensive two-dimensional gas-chromatographic mass-spectral analyses (GC x GC-MS) were carried out using a Pegasus 4D (Leco Corp., ST. Joseph, MI) equipped with a Restek Rxi-5 Sil MS primary column (5% diphenyl/95% dimethyl polysiloxane; 30m length x 0.25 mm i.d. x 0.25 -μm film thickness). Two different secondary columns were used during the course of this work. A Restek Rxi-5 Sil MS secondary column (5% diphenyl/95% dimethyl polysiloxane; 1.3 m length x 0.18 mm i.d. x 0.18 -μm film thickness) was used for chromatograms shown in Figure 1 and Figures S3 and S4 in the supplementary information. An SGE Analytical Science BPX50 secondary column (50% diphenyl/50% dimethyl polysiloxane; 1.0 m length x 0.1 mm i.d. x 0.10 -μm film thickness) was used for the chromatograms shown in Figure S5 in the supplementary information. An injector temperature of 250 °C with splitless injection was used, with the split valve opening at 20 seconds. The primary column oven temperature was held at 40 °C for 1 min, then increased at a rate of 3 °C/min to a temperature of 126°C, after which it was increased at a rate of 12 °C/min to a final temperature of 275°C. The secondary column oven followed a temperature program that maintained a +15°C offset from the primary column oven. A 3.5 s modulation period (0.80 hot pulse; 0.95 cold pulse) was used with a modulator temperature program that maintained a +15 °C offset from the secondary column oven. The carrier gas flow rate was 1.3 mL/min. Data processing was carried out using LECO ChromaTOF software with a signal-to-noise threshold of 30 for peak detection.

### GC-MS analyses

Additional analyses, including quantitative measurements, were carried out a Shimadzu QP2010 plus gas chromatograph mass spectrometer equipped with a Restek Rxi-5Sil MS column (5% diphenyl/95% dimethyl polysiloxane; 30m length x 0.25 mm i.d. x 0.25 -μm film thickness). An injector temperature of 250 °C with splitless injection was used, with the split valve opening at 0.52 minutes. A column temperature program was used that remained at 50 oC for the first minute, increased at a rate of 6 °C per min to a temperature of 200 °C, and then increased at a rate of 20 °C per min to a final temperature of 280 °C. The carrier gas flow rate was 1.22 mL/min.

### Chemotaxis assays in *C. elegans* and *C. remanei*

Chemotaxis assays were performed to evaluate the attractiveness of either worm-conditioned media (WCM) or prepared solutions of authentic methyl 3-methyl-2-butenoate (MMB). Two types of chemotaxis assays were conducted:

#### (1) WCM-based chemotaxis assays

These assays tested whether adult females of *C. remanei* or hermaphrodites of *C. elegans* are attracted to WCM derived from adult males of either species. The worm-conditioned media (WCM) (20 worms/10 µL) containing the native sex pheromone from *C. elegans* males and *C. remanei* males in M9 buffer was prepared and used for the chemotaxis assay according to the method in Leighton et al.,^10^ and applied to the inside lid of a 6-cm Petri dish prepared with chemotaxis media (2.0% agar, 1 mM MgSO_4_, 1 mM CaCl_2_, 25 mM KH_2_PO_4_, pH 6.0). A 10 µL aliquot of WCM was pipetted above one sodium azide spot, and 10 µL of M9 buffer was pipetted to the opposite side as a control (Fig. S1). The plate was then placed in a small box to eliminate the influence of light. The assay was run at least for 1 hr. If >20% of nematodes were still alive after 1 hr, we extended the incubation time until >80% of nematodes tested were dead, as determined by periodic checking. This took between 1 hr and 4 hrs. Pairwise comparisons were performed using a two-tailed Student’s t-test, with p < 0.05 considered statistically significant.

#### (2) MMB-based chemotaxis assays

These assays tested the attraction of *C. remanei* females to various concentrations of synthetic MMB. A dilution series ranging in concentration from 3.8 × 10^-4^ M to 3.8 × 10^-9^ M was prepared from a 50% (v/v) stock solution of MMB in ethanol. A 100 µL aliquot of each dilution was placed in the inverted cap of a microcentrifuge tube and positioned on the inside lid of a 10-cm Petri dish above one sodium azide spot (Fig. S2). Approximately 100 synchronized *C. remanei* females were placed at the center of the plate. The plate was then placed in a small box to eliminate the influence of light. The assay was run for at least 1 hr. If >20% of nematodes were still alive after 1 hr, we extended the incubation time until >80% of nematodes tested were dead, as determined by periodic checking. This took between 1 hr and 4 hrs. Differences among concentrations were analyzed using one-way ANOVA followed by Tukey’s multiple comparison test (p < 0.01).

In both types of assays, worms were bleached and synchronized, and L4 stage males/hermaphrodites/females were manually picked to ensure sexual purity. Worms were allowed to mature for an additional 24 hours before use. Worms were washed twice in M9 buffer and twice in ddH_2_O before being transferred to assay plates.

The chemotaxis index (CI) was calculated as {(number of nematodes at test cue zone) - (number of nematodes at the control zone)} /{(number of nematodes at test cue zone) + (number of nematodes at the control zone)}. All chemotaxis assays were repeated at least three times.

## Acknowledgments

Dr. Michael Milligan provided invaluable assistance with GC x GC-TOFMS analyses. We also thank Miwako Kobayashi for her assistance with chemotaxis assays.

## Author contributions

Conceptualization: YT, MG, XW, DL, FCS, PWS, RS Investigation: YT, MG, XW, MS, RS

Funding acquisition: FCS, PWS, RS Project administration: MG, PWS, RS

Supervision: PWS, RS

Writing – original draft: MG, XW, RS

Writing – review & editing: YT, MG, FCS, PWS, RS

## Data and materials availability

All data are available in the manuscript or the supplementary materials.

