## Supplementary Information for "A male-derived volatile sex pheromone in *Caenorhabditis* nematodes identified through its mimicry by a predator"

### Supplementary Figures

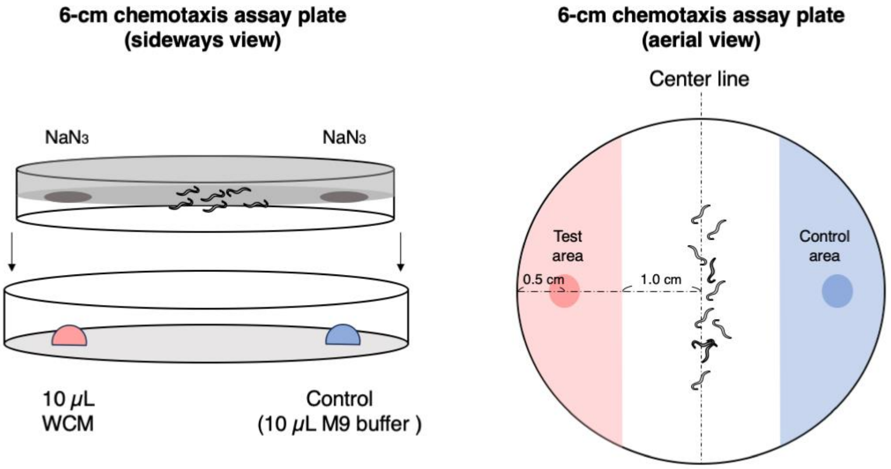


**Fig. S1: Schematic diagram of chemotaxis assay with worm conditioned media (WCM) on the 6-cm chemotaxis assay plate.**

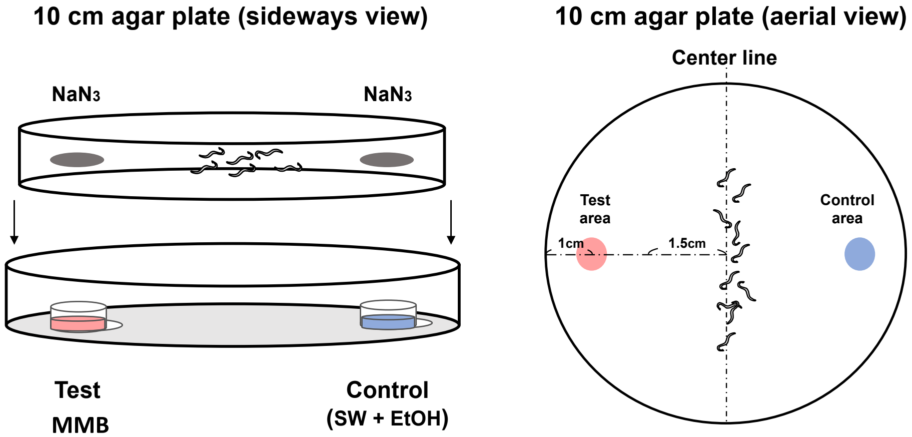


**Fig. S2: Schematic diagram of chemotaxis assay with MMB on the 10-cm chemotaxis assay plate. .**

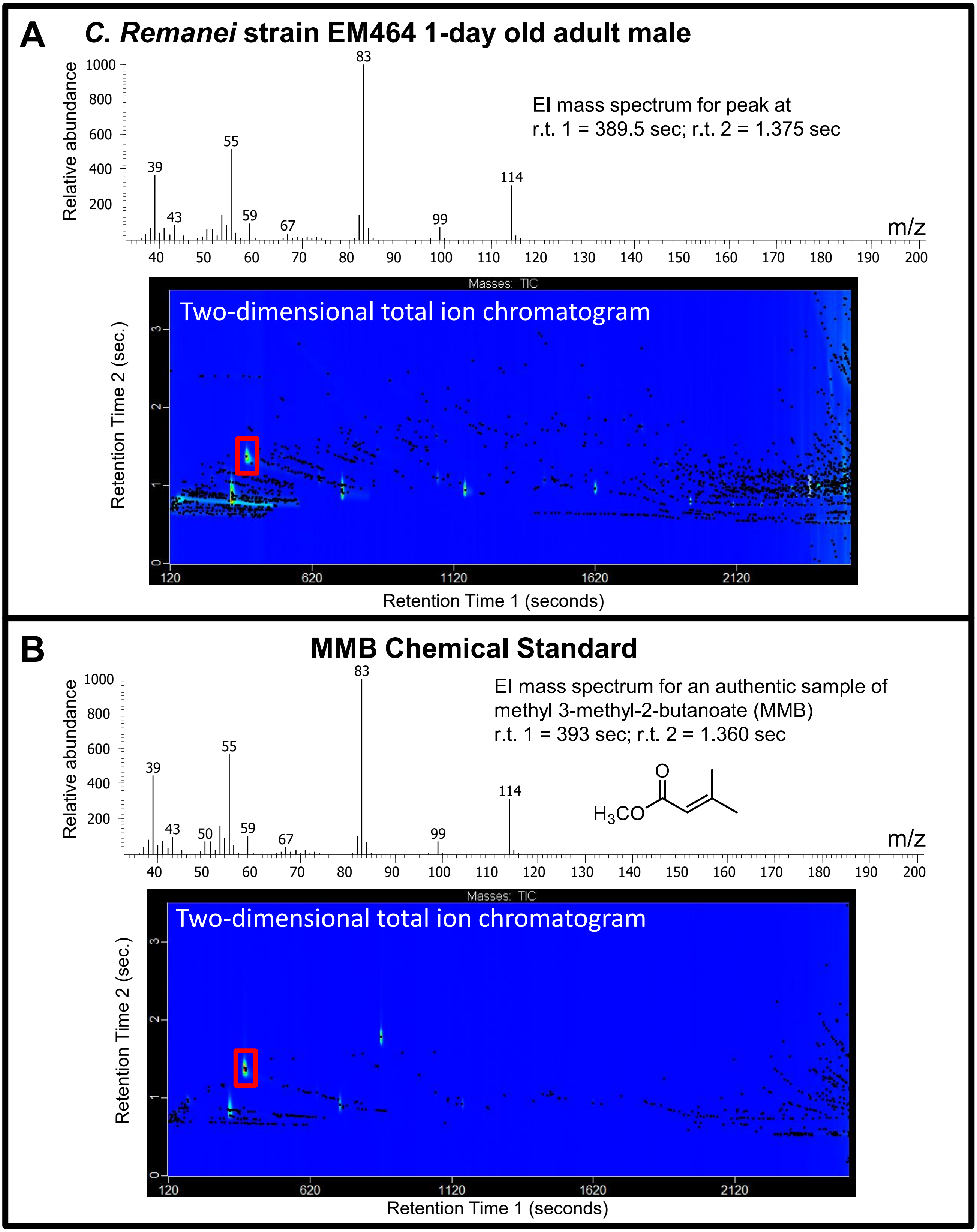


**Fig. S3: SPME headspace volatile sampling followed by GC x GC – TOFMS analyses revealed that MMB is produced by *C. remanei* adult males. (A)** EI mass spectrum and chromatogram (contour view) showing a chromatographic peak (indicated by red rectangle) which was unique to *C. remanei* adult male samples **(B)** Mass spectrum and chromatogram acquired for an authentic sample of MMB. Black dots represent discrete elution peaks identified during data processing (shown for a signal-to-noise threshold of 30).

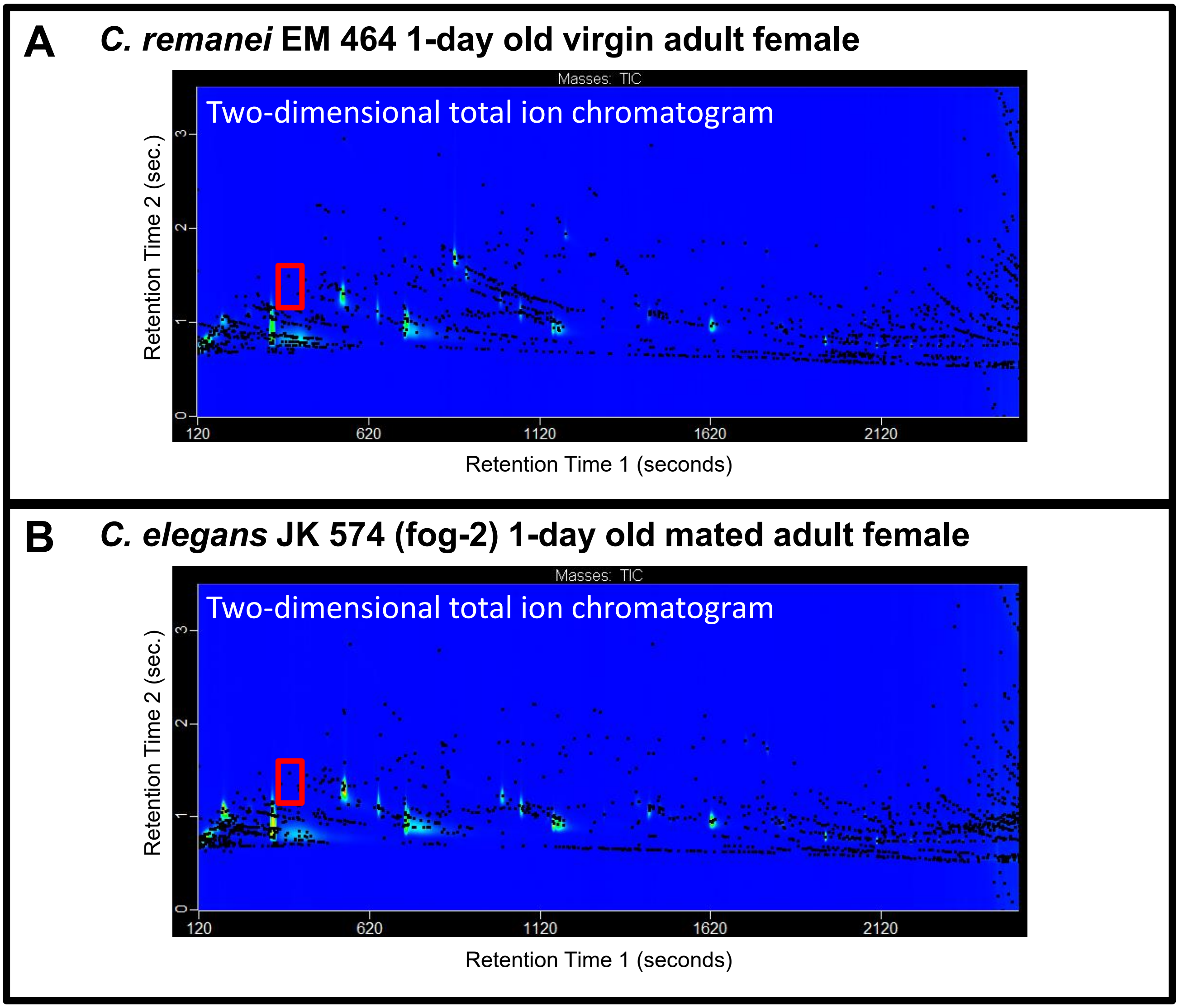


**Fig. S4** MMB was not detected for SMPE samples collected for *C. remanei* 1-day old virgin adult females, nor for any *C. elegans* sample. The red rectangle indicates the retention time region within which MMB would elute, if present.

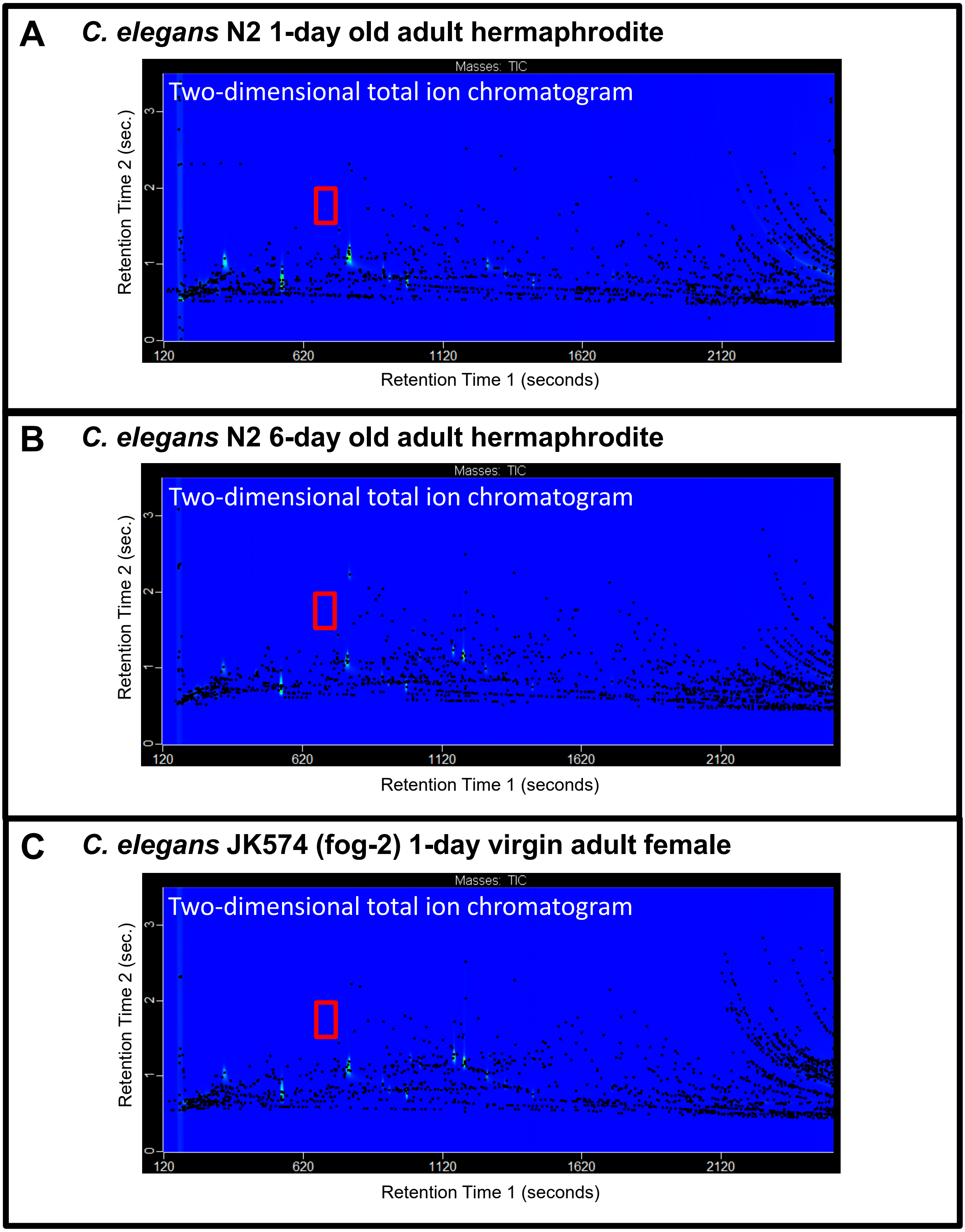


**Fig. S5** MMB was not detected for SMPE samples collected for any *C. elegans* sample. The chromatograms shown in this figure were acquired using an alternate secondary column (SGE Analytical Science BPX 50; described in Methods section), resulting in a change in the retention time region within which MMB would elute, if present (indicated by red rectangle).

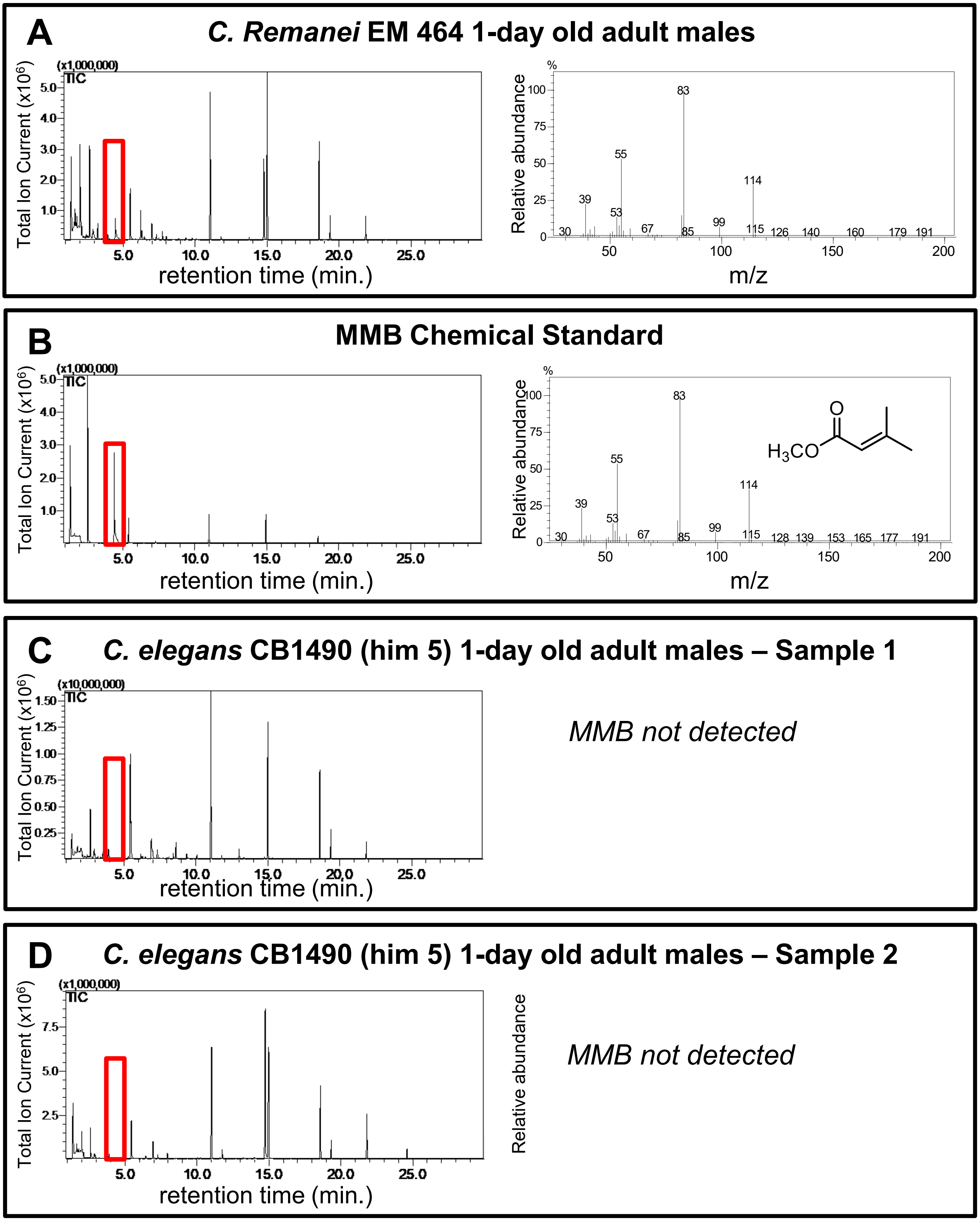


**Fig. S6 Additional analyses by SPME-GC-MS confirmed the presence of MMB for *C. remanei* adult male samples, as well as the absence of MMB for *C. elegans* adult males.** The red rectangle shows the retention time region within which MMB elutes, if present.
